# In silico pipeline for identification of targets of plastic-derived chemicals

**DOI:** 10.64898/2026.11.22.768400

**Authors:** Alan Gabriel Halatian, Luciana Rodriguez Sawicki, Leandro Matías Sommese, Marina Olga Fernandez, Ana Julia Velez Rueda

## Abstract

Plastic pollution is a growing global health concern, yet the specific human protein targets mediating the toxic effects of plastic-derived chemicals (PDCs) remain largely unknown. Here, we present an integrative in silico pipeline that combines transcriptomic data with structural bioinformatics to identify putative human plastic-derived targets. Through comparative analysis with their canonical ligands, we identified 27 potential targets showing global binding energies comparable to endogenous ligands (-5.26 vs -5.60 kcal/mol), and six cases in which plastic-derived compounds performed better. Chemical similarity analysis revealed very low structural resemblance between PDCs and canonical ligands, indicating that PDCs do not act as structural mimetics but instead exploit different interactions within the same binding cavities.

In a proof-of-concept case, we showed that Deoxyribonuclease I (DNase I) binds BP2 with higher affinity than its canonical ligand PO₄-^3^ (median -5.85 vs -2.55 kcal/mol). It also forms hydrogen bonds with Asn192 and π-stacking interactions with the catalytic histidine His274, a pattern conserved in bovine DNase I. Experimental validation confirmed that BP2 at 10^-⁵^ M inhibits DNA hydrolysis by 21.5% in bovine DNase I (p-value < 0.01) and 23.2% in recombinant human DNase I (p-value < 0.001). Our work establishes a robust, transferable computational framework for identifying protein targets of environmental contaminants and reveals that PDCs engage their targets through novel non-mimetic binding modes, with structural patterns that can be conserved across species. These findings may have significant implications for the safety assessment of plastic additives.

**Highlights:**

- An integrative in silico pipeline identifies putative human targets of plastic-derived chemicals (PDCs) by combining transcriptomic data with structural bioinformatics.
- PDCs exhibit global binding energies comparable to canonical ligands and exceed them in six cases, without acting as structural mimetics.
- BP2 binds DNase I with higher affinity than its canonical ligand PO₄³⁻ and engages in π-stacking with the catalytic histidine His274.
- The BP2–DNase I interaction pattern is conserved in bovine DNase I, indicating cross-species relevance.
- Experimental validation confirms that BP2 inhibits DNA hydrolysis by 21.5% in bovine DNase I and 23.2% in human DNase I.

## Introduction

Plastics have transformed our daily lives, and have brought significant economic benefits to the industrial sector. For this reason, their use and annual production continue to grow steadily, even when their environmental impact is undeniable ^1–3^. It is estimated that by 2050 plastic use will reach 884 million tonnes, up from 464 million tonnes recorded in 2020 ^4^. This increasing use is accompanied by plastic accumulation that can neither be eliminated nor recycled, and is estimated to reach 4,725 million tonnes by the same year ^5^. Current valuations suggest that the amount of plastic pollution entering terrestrial and aquatic environments will increase by 710 million tonnes by 2040, even if immediate action is taken to reduce waste ^6^.

In the environment, plastics generally degrade and fragment due to various processes, including bacterial activity, UV exposure, temperature fluctuations, and abrasion ^7^. Additionally, polymers are mixed with chemical additives that have been described as toxic, such as phthalate plasticisers, brominated flame retardants, and antimicrobial agents ^8,9^, among others, which are capable of continuously leaching from their source materials into food, water and soil. This generates universal human exposure that is detectable in the vast majority of the general population, throughout the whole life cycle of plastics ^10,11^.

Plastics and their additives contaminate soils and watercourses, are ingested and incorporated by production animals, and reach humans through the diet, air and drinking water ^12–14^. In this sense, human, animal, and ecosystem health are expressions of a single system under stress induced by pollution ^15,16^. In the context of environmental stress, plastic derivatives can act as disruptors of the native protein state, either by generating reactive oxygen species that damage proteins, altering the pH of cellular compartments, or directly interfering with the folding and action of specific proteins ^16^.

At the molecular level, pollutants can induce misfolding or trap proteins in an inactive conformation, triggering cellular responses that seek to compensate for the loss of function and restore protein homeostasis ^17,18^. These responses involve activating signalling pathways that reprogram gene expression to increase the production of chaperones, folding factors and degradation systems, following the conformational sequestration of one protein and a compensatory increase in the expression of others ^19^.

In addition, chronic activation of these pathways can exceed an adaptive threshold, generating cellular accumulations that not only increase ligand-protein interactions but can also lead to apoptosis, thereby contributing to the cellular toxicity observed with certain plastics ^1,2,18^. Evolution has generated quality control mechanisms and cellular stress responses that continuously monitor the conformational landscape and actively compensate for deviations from protein homeostasis.

In the context of environmental stress induced by plastic pollutants, identifying the molecular targets affected by their derived compounds is crucial to understanding the impact on human and environmental health. Although the general toxic effects of plastics are known, the molecular basis of their interaction with specific proteins and their potential interference with endogenous ligands has not yet been thoroughly explored. In this work, we propose an in silico analysis pipeline that integrates experimental data, structural modelling and molecular docking to identify potential protein targets of plastic-derived compounds (PDCs), evaluating their ability to compete with canonical ligands and disrupt key biological functions. This approach also allows us to extrapolate our findings to species that are part of the human food chain, laying the groundwork for future experimental validation and risk assessment.

## Results

### Global ligand-target binding energies comparison

An initial list of 12 compounds was selected from the literature on plastic additives and environmental contaminants to cover the main chemical classes of plastic-derived endocrine disruptors ^7,11,19,20^. These comprised nine phthalates: bis(2-ethylhexyl) phthalate (DEHP), diisoheptyl phthalate (DIHP), diisohexyl phthalate (DIHXP), diundecyl phthalate (DUP), diisononyl phthalate (DINP), diisooctyl phthalate (DIOP), dipropyl phthalate (DPP), ditridecyl phthalate (DTDP), and butyl benzyl phthalate (BBP); and three phenolic compounds: bisphenol A (BPA), benzophenone-2 (BP2), and benzophenone-3 (BP3). Afterwards, this number was reduced to 8 representative PDCs by structural clustering using the Butina algorithm ^21^: five phthalates (BBP, DPP, DEHP, DIOP, DTDP) and three phenolic compounds (BP2, BP3, BPA).

Physicochemical characterization (Table 1) revealed that the phenolic compounds (BP2, BP3, BPA) have the lowest fraction of sp^3^-hybridized carbons (Fsp^3^ = 0.0–0.20), indicating planar, rigid structures favorable for π-type interactions. BP2 stands out with the highest polar surface area (TPSA = 97.99 Å²) and hydrogen-bonding capacity (HBD = 4, HBA = 5). In contrast, the phthalates show increasing conformational flexibility and lipophilicity with chain length, reaching extremes in DTDP (LogP = 10.62; 26 rotatable bonds). All phthalates lack hydrogen-bond donors (HBD = 0), predicting hydrophobic binding modes. The Quantitative Estimate of Drug-likeness (QED) was highest for the smaller bisphenols, such as BPA and BP3 (0.83 and 0.82), indicating greater suitability for protein binding than the larger phthalates, such as DTDP (0.09). These physicochemical differences suggest that these compounds may establish a wide diversity of possible interaction types with human protein targets.

**Table 1.** Physicochemical descriptors of the eight selected PDCs, calculated with RDKit. Family: chemical classification according to the predominant functional group. Molecular weight: molecular mass in Daltons. LogP: octanol–water partition coefficient, an indicator of lipophilicity. TPSA: topological polar surface area (Å²), related to permeability and hydrogen-bonding capacity. H donors / H acceptors: number of hydrogen-bond donors and acceptors. Aromatic rings: number of aromatic systems in the molecule. Fsp³ fraction: proportion of sp³-hybridized carbons relative to total carbons, an indicator of the three-dimensional character and saturation of the molecule. QED composite score from 0 to 1, integrates multiple physicochemical properties as a global indicator of molecular suitability for protein binding, indicating that the smaller bisphenols are more suitable for protein binding than the larger, more lipophilic phthalates.

| PDC | Molecular Weight (Da) | Log P | TPSA (Å<sup>2</sup>) | H Donors | H Acceptors | Aromatic Rings | Fsp<sup>3</sup> | QED |
|---|---|---|---|---|---|---|---|---|
| BP3 | 228.25 | 2.63 | 46.53 | 1 | 3 | 2 | 0.07 | 0.82 |
| BP2 | 246.22 | 1.74 | 97.99 | 4 | 5 | 2 | 0 | 0.6 |
| BPA | 228.29 | 3.42 | 40.46 | 2 | 2 | 2 | 0.2 | 0.83 |
| BBP | 312.36 | 4 | 52.6 | 0 | 4 | 2 | 0.26 | 0.57 |
| DEHP | 390.56 | 6.43 | 52.6 | 0 | 4 | 1 | 0.67 | 0.34 |
| DTDP | 530.83 | 10.62 | 52.6 | 0 | 4 | 1 | 0.76 | 0.09 |
| DPP | 250.29 | 2.82 | 52.6 | 0 | 4 | 1 | 0.43 | 0.73 |
| DiOP | 390.56 | 6.43 | 52.6 | 0 | 4 | 1 | 0.67 | 0.27 |

While the molecular bases of PDCs toxic effects in zebrafish have been extensively studied ^22,23^, identifying the specific human protein targets that mediate effects of PDCs remains poorly explored and constitutes a challenge. To address this, we started from 326 plastic-responsive genes described in Lomonta’s work that provided a detailed multi-level analysis, used for identifying human orthology. From the resultant list of human orthologs (159 total), we focused on 27 proteins having a characterized canonical ligand. Since the experimental PDB structures were available for only 27.7% of these proteins, but were unsuitable because of missing residues (12% on average), we used AlphaFold models for all 27 proteins: 25 had a pLDDT above 80, one had 74 and one had 65.

All eight PDCs and the canonical ligands were docked against the 27 putative targets. The global binding energy distributions observed for both groups of ligands were similar, with median binding energies of −5.26 kcal/mol (95% CI −5.38 to −5.08; n = 472) for PDCs and −5.60 kcal/mol (95% CI −5.86 to −5.27; n = 141) for canonical ligands, suggesting PDCs could be able to compete with endogenous ligands for binding. In 18.6% of the cases, at least one PDC performed better than the canonical ligand, while in only 17% of cases the canonical ligand was favored. In the other cases, differences were below the error threshold (2.85 kcal/mol), indicating comparable affinities. Chemical similarity analysis using Morgan fingerprints ^24^ revealed very low structural similarity between PDCs and their corresponding canonical ligands (median Tanimoto coefficient = 0.054; maximum = 0.136). This value was even lower (0.033) when PDCs showed better binding energy, compared to 0.060 when canonical ligands were favored. These results may indicate PDCs do not act as structural mimetics of endogenous ligands, even when they can compete with canonical ligands. No correlation was found between differential affinity and cavity descriptors (volume, polarity, hydrophobicity).

Analysis of all 30 docking comparisons in which PDCs outperformed their corresponding canonical ligands revealed that PDC binding is primarily driven by π interactions and hydrophobic contacts (Figure 1). This is consistent with the chemical nature of the PDCs: the phenolic compounds (BP2, BP3, BPA) contribute planar aromatic rings capable of π–π stacking, while the phthalates contribute long, flexible alkyl chains that engage in hydrophobic contacts. The high frequency of Glu (Figure 1) likely reflects its abundance in the binding pockets of the 27 target proteins, many of which are hydrolases and GTPases with polar catalytic residues. However, because Glu cannot engage in π-stacking, its prevalence also suggests that polar contacts (e.g.: hydrogen bonds and electrostatic interactions) contribute to PDC binding alongside the dominant hydrophobic and π-type interactions. In contrast, aromatic residues such as Trp and Tyr, though less frequent overall (Figure 1), may contribute disproportionately through π-stacking interactions. Together, these findings indicate that PDCs bind through mechanisms that exploit the chemical features of both the ligand and the protein binding site.

**Figure 1.**
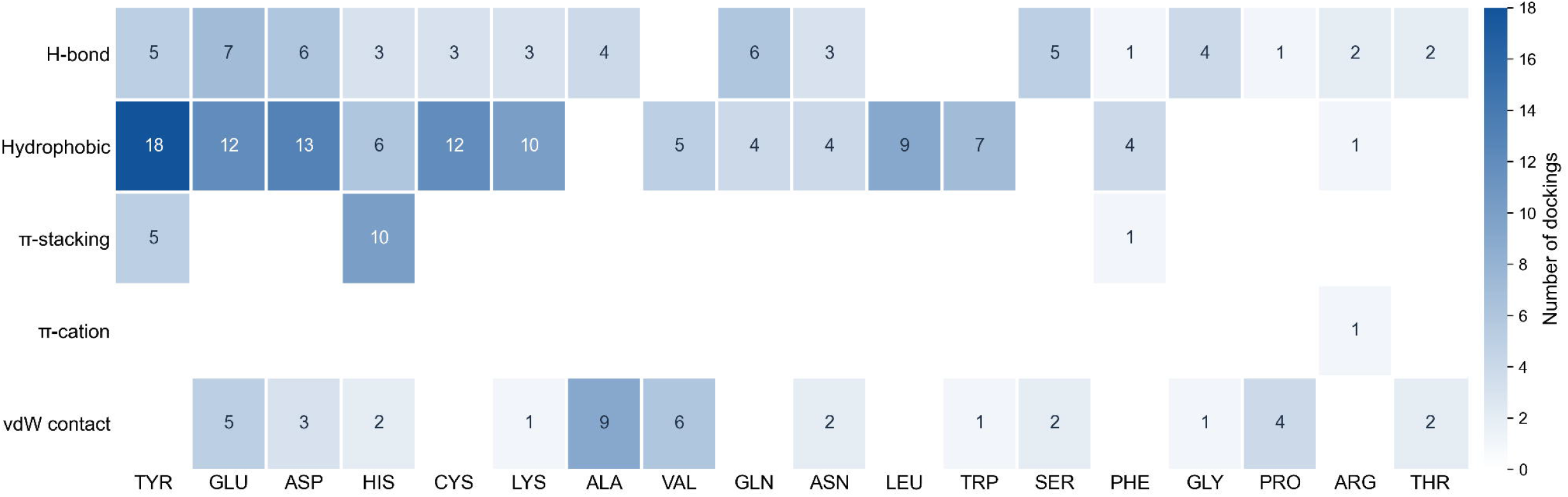
Residue–interaction pairs exclusive to the PDC in the dockings where it presented better binding energy. Number of dockings in which the PDC forms each type of interaction (rows) with each amino-acid type (columns) and the canonical ligand in the same pocket does not. Residues are grouped by type and sorted by total count.

### Functional characterization of candidate proteins

Regarding biological processes, the most enriched term among proteins with preferential affinity for PDCs was the immune system, followed by programmed cell death and extracellular matrix organization, consistently with the known toxicological profiles of phthalates and bisphenols. Notably, the subset of proteins with higher affinity for the canonical ligand showed significant enrichment in lipid metabolic processes, as well as in terms related to anatomical structure development and nervous system development –a trend likely reflecting the apolar nature of these endogenous ligands and their specific physicochemical binding requirements. Other processes, such as cell motility and reproductive processes, showed a balanced distribution between both types of ligands.

The analysis of molecular functions revealed that hydrolase activity was the most enriched term, with a predominance of proteins in which the PDC presented better binding energy. A higher proportion of proteins with affinity for PDCs was also observed in functions such as GTPase activity, molecular function regulator activity, and molecular sequestering activity. In contrast, oxidoreductase activity, isomerase activity, and transferase activity were predominantly associated with proteins in which the canonical ligand was the winner. The predominance of PDCs in proteins with hydrolase and GTPase activity is particularly relevant, given that these enzymes participate in intracellular signaling cascades and in the processing of essential substrates for cellular homeostasis. This functional profile suggests that PDCs may interfere with key regulatory nodes in cell signaling and metabolism, potentially through non-competitive binding modes that exploit hydrophobic and π-type interactions, as described in the previous section.

The KEGG disease enrichment analysis of proteins with preferential PDC binding revealed significant associations with systemic lupus erythematosus (H00080), chronic myeloid leukemia (H00004), and hemochromatosis (H00211), among others (Supplementary Table 1). The SLE association is of particular interest given that Deoxyribonuclease I (DNase I), a protein with strong PDC preference in our docking analysis, is critical for the clearance of neutrophil extracellular trap (NET) and its deficiency is a known SLE risk factor ^25^. Moreover, other PDCs have been reported to induce NETosis ^26,27^, suggesting a potential mechanistic link between PDC exposure and autoimmune pathology that warrants experimental investigation. The enrichment of cancer-related diseases (CML, B-ALL) aligns with the presence of ABL1 and MAPK12 in our target set, both of which are implicated in oncogenic signaling and cell survival^28^. The convergence of our computational predictions with established disease associations and the reported toxicological effects of plastic additives strengthens the biological plausibility of our findings and provides us with a prioritized list of targets for future experimental validation. These findings also underscore the value of this integrative approach for identifying environmental health risks.

### Case of biological relevance: Deoxyribonuclease I

To illustrate the structural and functional relevance of the identified targets, we selected DNase I as a representative candidate from the 27 identified proteins. The selection was guided by three complementary lines of evidence: (i) it showed consistent preferential binding affinity for multiple PDCs in our docking analysis (particularly BP2, BP3, and BPA); (ii) it was overexpressed in the zebrafish transcriptomic data from Limonta et al.^22^, which may indicate a compensatory response to functional impairment induced by pollutant exposure ^29–31^; and (iii) biologically, in humans, it has been shown that plays fundamental roles in DNA degradation during apoptosis and in the NETs clearance; its deficiency has been associated with the development of SLE (H00080) ^25^.

DNase I is ubiquitously expressed but primarily secreted by the exocrine pancreas. Its function relies on the nonspecific hydrolysis of phosphodiester bonds in DNA, requiring Ca²⁺ for electrostatic optimization and Mg²⁺ for catalysis. Across an evolutionary alignment of 206 orthologous sequences from 153 vertebrate and non-vertebrate chordate species, DNase I exhibited strong overall conservation, with a mean pairwise sequence similarity of 70.5% (55.8% identity), and particularly high conservation at the metal-coordinating residues ^32^. DNase I is conserved across the 206 sequences of the ortholog alignment (153 species), with a mean pairwise sequence similarity of 70.5% (55.8% identity). The Ca²⁺-binding site showed strong conservation at Asp223 (99.0%) and Thr225 (98.5%) but high variability at Thr227 (35.0%) (Figure 2C). The Mg²⁺-binding site was conserved at Asn29 (97.1%) and Glu61 (97.6%) and moderately conserved at Asp273 (81.1%) (Figure 2D). Catalytic residues ranged from highly conserved (Asp190, Asn192, and His156; 98.0% identity) to moderately variable (His274; 77.7%), though all retained functional relevance based on mutagenesis data (Figure 2B). Similarly, the DNA-binding site exhibited heterogeneous conservation, with highly conserved core contact residues (95.0%) and variable peripheral positions such as Pro228 (1.94%) (Figure 2E). Collectively, these results underscore the structural robustness of catalytically essential regions while suggesting peripheral variability that could enable species-specific or environmental adaptations.

**Figure 2.**
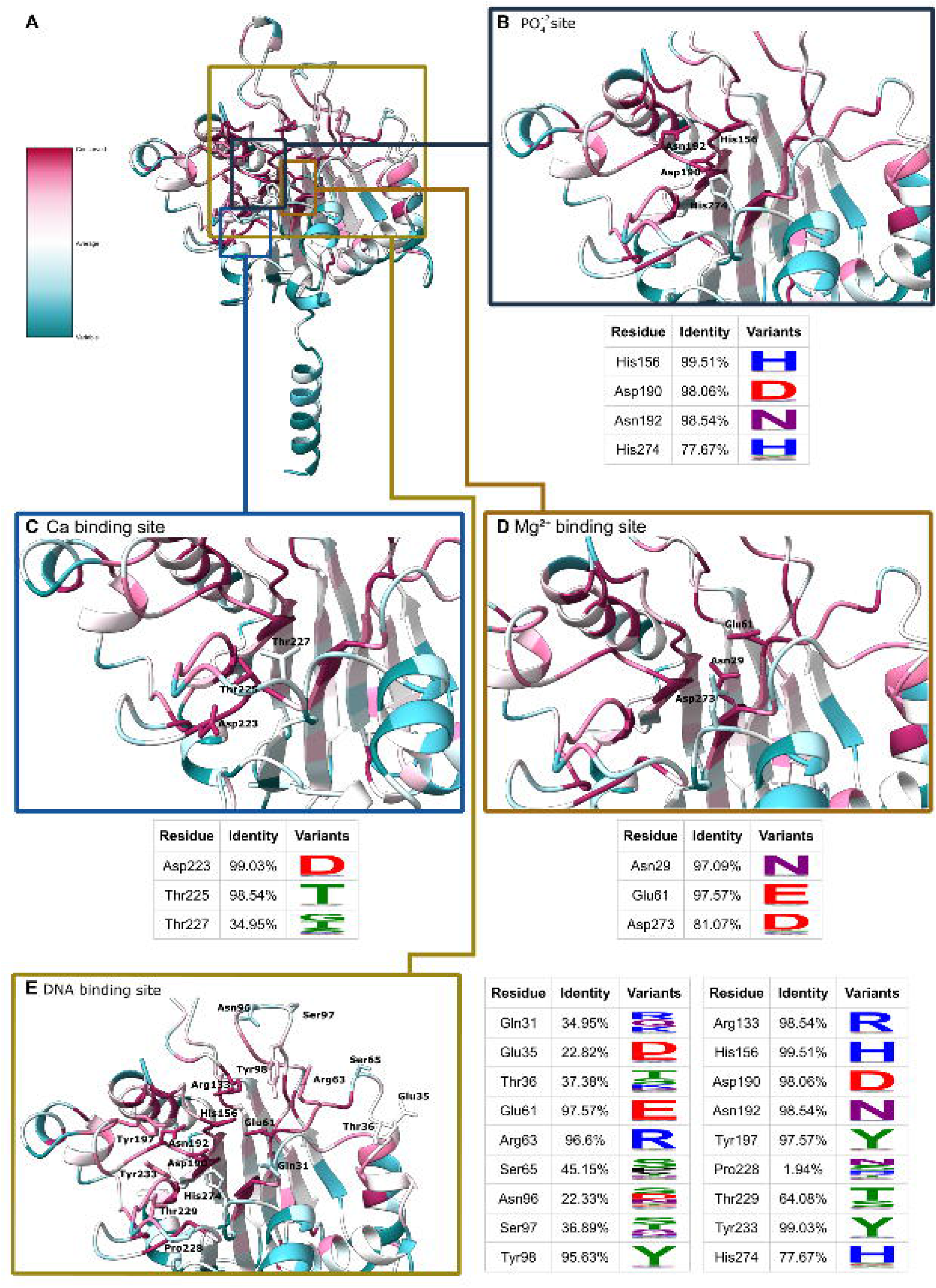
Conservation of the functional sites of human DNase. **I.** (A) Structure of human DNase I colored by residue conservation across 206 orthologous sequences (pink, conserved; blue, variable). Colored boxes mark the binding sites shown in detail in B–E. (B) PO₄²⁻, (C) Ca²⁺, (D) Mg²⁺ and (E) DNA binding sites. The table accompanying each site gives, for each residue, the percentage of sequences with the human residue at that position (Identity) and a sequence logo of the residues found there (Variants).

The cavity selected for analysis of the interaction with PDCs was the one showing high coincidence with the active site annotated in UniProt for human DNase I (UniProt ID P24855). The predicted cavity includes the functional regions previously described: the two residues that coordinate the catalytic Mg²⁺ (Glu61, which coordinates directly, and Asp273, which does so through a water-mediated interaction); the catalytic core that acts on the scissile phosphate (His156, Asp190, Asn192, His274); three direct DNA-binding residues reported in the bovine complex (Tyr197, Thr229, Tyr233); and two residues of the adjacent Ca²⁺ site (Thr225, Thr227).

The canonical ligand, PO₄³⁻, which was annotated and used for comparisons, showed binding energies of around −2.55 kcal/mol, markedly weaker than the binding energies predicted for PDCs. BP2 showed the highest affinity (−6.21 to −5.47 kcal/mol; median −5.85), followed by BP3 (−5.95 to −4.88 kcal/mol; median −5.26) (Table 2). Notably, the observed affinity gradient followed the aromaticity and rigidity of the compounds rather than their polarity. The four PDCs bearing two aromatic rings (BP2, BP3, BBP, and BPA) occupied four of the five highest positions, whereas the flexible, aliphatic phthalates carrying a single aromatic ring (DPP, DIOP, and DTDP) displayed the lowest affinities. Ranking first, BP2 was the only compound combining a high polar surface area (97.99 Å²) with four hydrogen-bond donors.

**Table 2.** Binding affinities and interaction profiles of PDCs against human DNase I. Values represent median binding energies (kcal/mol) between conformers and the percentage of docking poses exhibiting each interaction type. Hydrogen bonds and π stacking emerge as the discriminating interactions, while VdW/hydrophobic contacts are ubiquitous across all compounds

| Compound |  | Median Binding Energy (Kcal/mol) | H-Bond Acceptor (%) | H-Bond Donor (%) | π Stacking (%) | VdW/Hydrophobic (%) |
|---|---|---|---|---|---|---|
| PDC | BP2 | -5.85 | 67 | 78 | 33 | 100 |
|  | BP3 | -5.26 | 44 | 44 | 56 | 100 |
|  | BPA | -4.88 | 56 | 44 | 56 | 100 |
|  | BBP | -5.03 | 33 | 11 | 33 | 100 |
|  | DEHP | -4.91 | 22 | 0 | 0 | 100 |
|  | DIOP | -4.32 | 11 | 0 | 0 | 100 |
|  | DPP | -4.03 | 11 | 0 | 22 | 100 |
|  | DTDP | -3.57 | 0 | 0 | 0 | 100 |
| Canonical | PO<sub>4</sub><sup>3-</sup> | -2.55 | 100 | 0 | 0 | 67 |

To understand the molecular determinants of these affinity differences, we analysed the protein-ligand interactions using fingerprints (Figure 3A). Van der Waals and hydrophobic contacts were present in 100% of the conformations of all evaluated PDCs, and therefore did not constitute discriminating factors between compounds. Instead, the interactions that explained the observed energetic differences were: hydrogen bonds and π stacking. The richest profile was observed for BP2, which formed hydrogen-bond acceptor and donor interactions in 67% and 78% of its conformations, respectively, together with π stacking in 33%, consistent with its highest predicted affinity. BP3 and BPA also showed diverse profiles: BP3 stood out for the high frequency of π stacking (56%), while BPA combined hydrogen-bond acceptor (56%), hydrogen-bond donor (44%), and π-stacking (56%) interactions. At the opposite extreme, DEHP and DIOP completely lacked hydrogen-bond donor and π-stacking interactions. These results may indicate that binding affinity is determined not by the quantity of contacts, but by their chemical specificity. Thus, PDCs enriched in polar interactions and π stacking achieve higher affinities, whereas profiles dominated by nonspecific contacts achieve weaker binding despite higher contact density.

**Figure 3.**
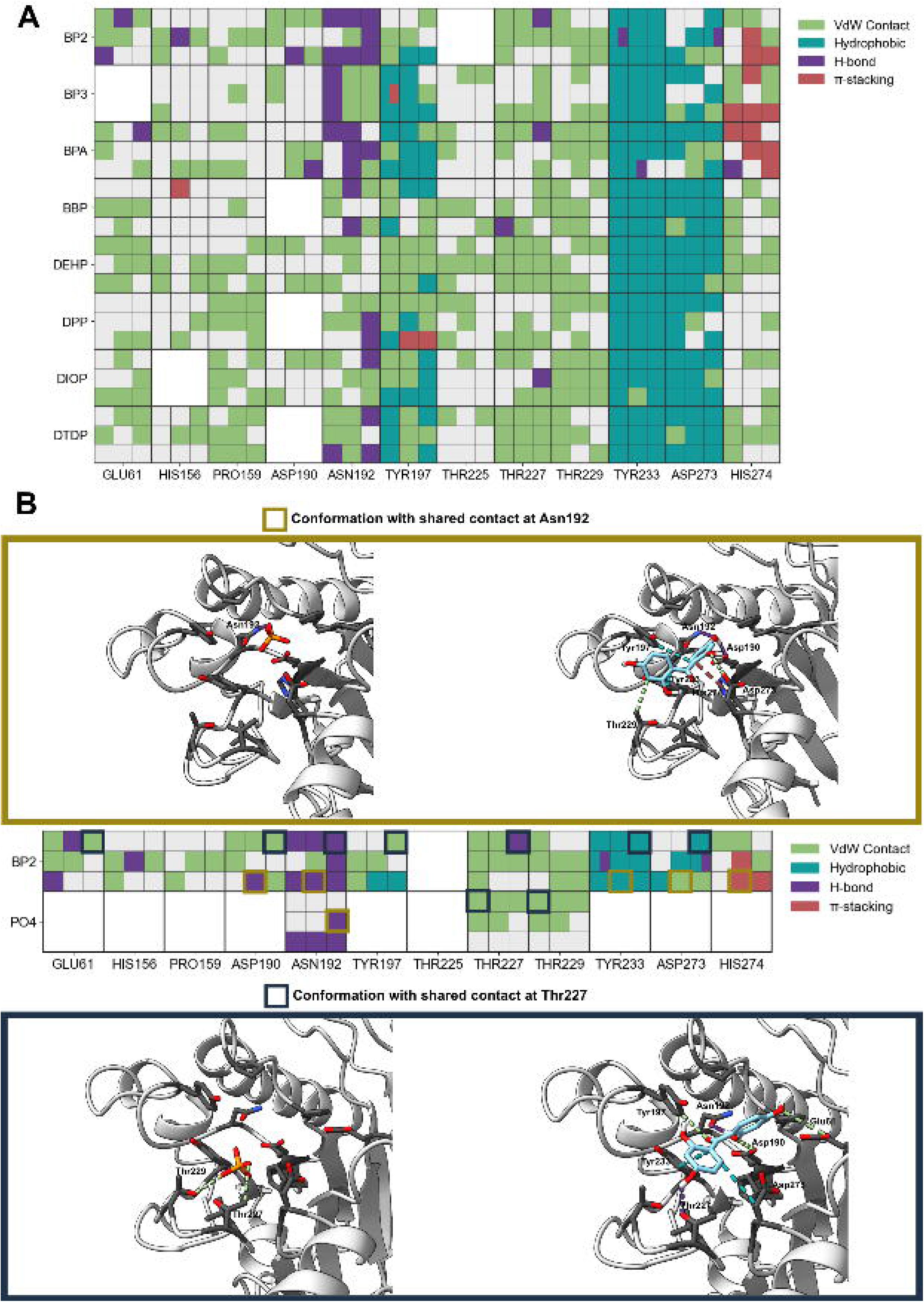
Protein-ligand interaction fingerprints of PDCs and comparison with the canonical ligand PO₄³⁻. (A) Interaction fingerprint matrix of the eight studied PDCs against the residues of the binding site. Rows correspond to the PDCs and columns to the evaluated residues. Each cell corresponds to a 3 × 3 submatrix, in which each element represents a molecular docking pose and the color indicates the type of interaction observed: van der Waals contact (green), hydrophobic interaction (teal), hydrogen bond (purple) and π-stacking (red). White cells indicate absence of contact in the corresponding pose. (B) Directed comparison between poses of BP2 and PO₄³⁻ that share contacts with residues of the binding site. Gold boxes indicate poses in which both ligands contact Asn192; the corresponding structural visualization is shown in the upper block. Dark blue boxes indicate poses in which both ligands contact Thr227; in this case, PO₄³⁻ additionally extends its interaction towards Thr229, as shown in the lower block. Contact residues are indicated with labels and interactions follow the same color code as in (A).

The comparison between the BP2 profile and the canonical ligand showed that PO₄³⁻ predominantly forms hydrogen bonds with Asn192 and van der Waals contacts with Thr227 and Thr229 (Figure 3B). Notably, these contacts do not coexist in a single conformation: hydrogen bonds with Asn192 and van der Waals interactions with the threonines correspond to distinct conformations occupying separate sectors of the cavity. In contrast, BP2 and BP3 replicate the hydrogen bond with Asn192 while additionally engaging in π-stacking with the catalytic histidine His274; BP3 further extends this interaction toward Tyr197. Both compounds complement these contacts with hydrophobic interactions involving Tyr197 and Tyr233, residues critical for DNA binding, as well as Asp273, which coordinates the catalytic Mg²⁺. The π-stacking between BP2 and His274 is particularly noteworthy, as it mirrors the interaction pattern reported for synthetic DNase I inhibitors, which establish π interactions and hydrogen bonds with the catalytic histidines of the active pocket ^33^. This interaction-level mimicry, rather than structural resemblance to the endogenous ligand, supports the hypothesis that BP2 disrupts catalytic activity.

### Cross-species impact analysis

The European Food Safety Authority (EFSA) has recognized the dietary route as the main route of exposure to plastic derivatives in the general population, and for this reason we focused on cattle (Bos taurus), a major source of meat for human consumption. Human and bovine DNase I are particularly close (87.9% similarity, 77.0% identity), and their structures superimpose with an RMSD of 0.32 Å over 260 Cα atoms, with catalytic residues and active-site geometry that remain practically identical (Figure 4A).

**Figure 4.**
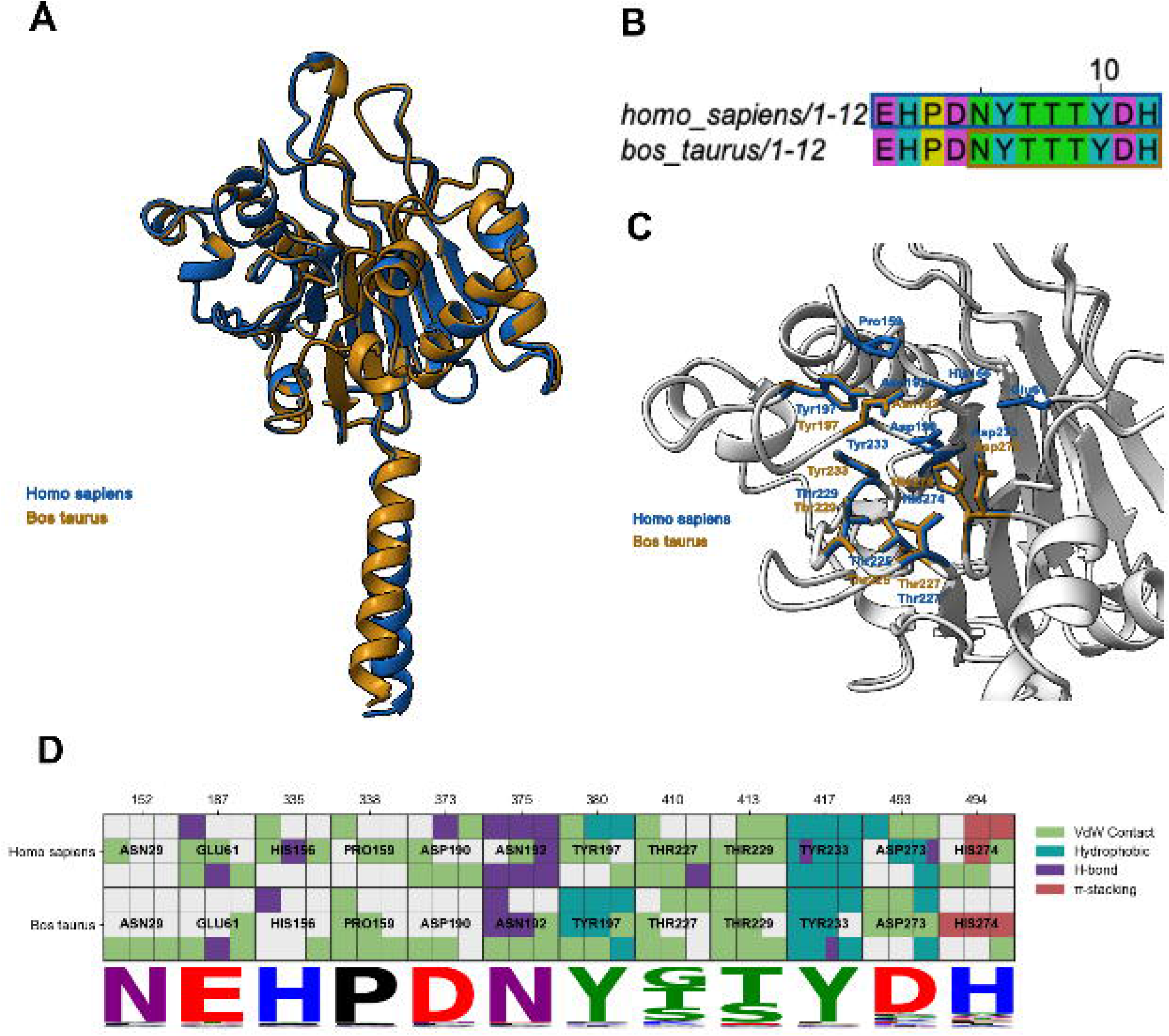
Conservation of the docking cavity between human and bovine DNase. **I.** (A) Structural superposition of human (blue) and bovine (brown) DNase I. (B) Alignment of the cavity residues. The boxes mark the residues that form the cavity in each species. (C) Superposition of the cavity residues of both species. (D) Interaction fingerprint of BP2 with the cavity residues of each species. Each cell holds the nine docking poses (3 × 3), colored by interaction type, with gray meaning no contact. The sequence logo below shows the conservation of each position across the alignment.

The molecular docking analysis previously performed for the human protein was extended to Bos taurus using the same PDC ligand, assigning the target cavity by homology through structural alignment. The predicted binding cavity in the bovine protein was considerably smaller than that in the human protein, with only 8 of the 12 residues present in the human site, and a smaller volume: 237 Å³ for the bovine protein compared with 316 Å³ for the human protein. Considering only the active site’s common residues, the RMSD observed in the structural alignment was 1.701 Å (Figure 4B and 4C).

Despite these differences, the docking analysis showed that the key interactions established by BP2 are conserved between human and Bos taurus. In both species, BP2 forms a hydrogen bond with Asn192 and engages in π-stacking with the catalytic histidine His274, along with hydrophobic contacts involving Tyr197, Tyr233, and Asp273. Notably, the hydrogen bond between Asn192 and the canonical ligand PO₄³⁻ is also preserved in Bos taurus, indicating that this position remains functionally relevant across species.

The binding energy landscape was also similar. Bovine DNase I showed a range of energies between -5.49 and -4.77 kcal/mol, with a median of -5.22 kcal/mol, comparable to the values obtained for the human protein. These results suggest that the structural determinants of BP2 binding are largely conserved within species, and support the hypothesis that the inhibitory effect may extend to this species.

To assess how BP2 affects DNA binding, we calculated for each residue the percentage of simulation frames in which it contacted the DNA (occupancy) (Figure 5). In the absence of BP2, DNA binding was primarily mediated by residues previously identified in DNase I–DNA crystal structures. In human DNase I, these residues were Thr36 (95%), Arg133 (83%), Glu61 (82%), Tyr98 (67%), Arg63 (60%), and the catalytic His156 (58%), whereas in bovine DNase I, they were Arg31 (91%), Thr36 (81%), His156 (52%), Arg63 (46%), and Tyr197 (42%). BP2 did not abolish DNA binding; in both species, the DNA remained in contact with the protein throughout the simulation, and Thr36 retained near-complete occupancy (99% in human and 100% in bovine DNase I).

**Figure 5.**
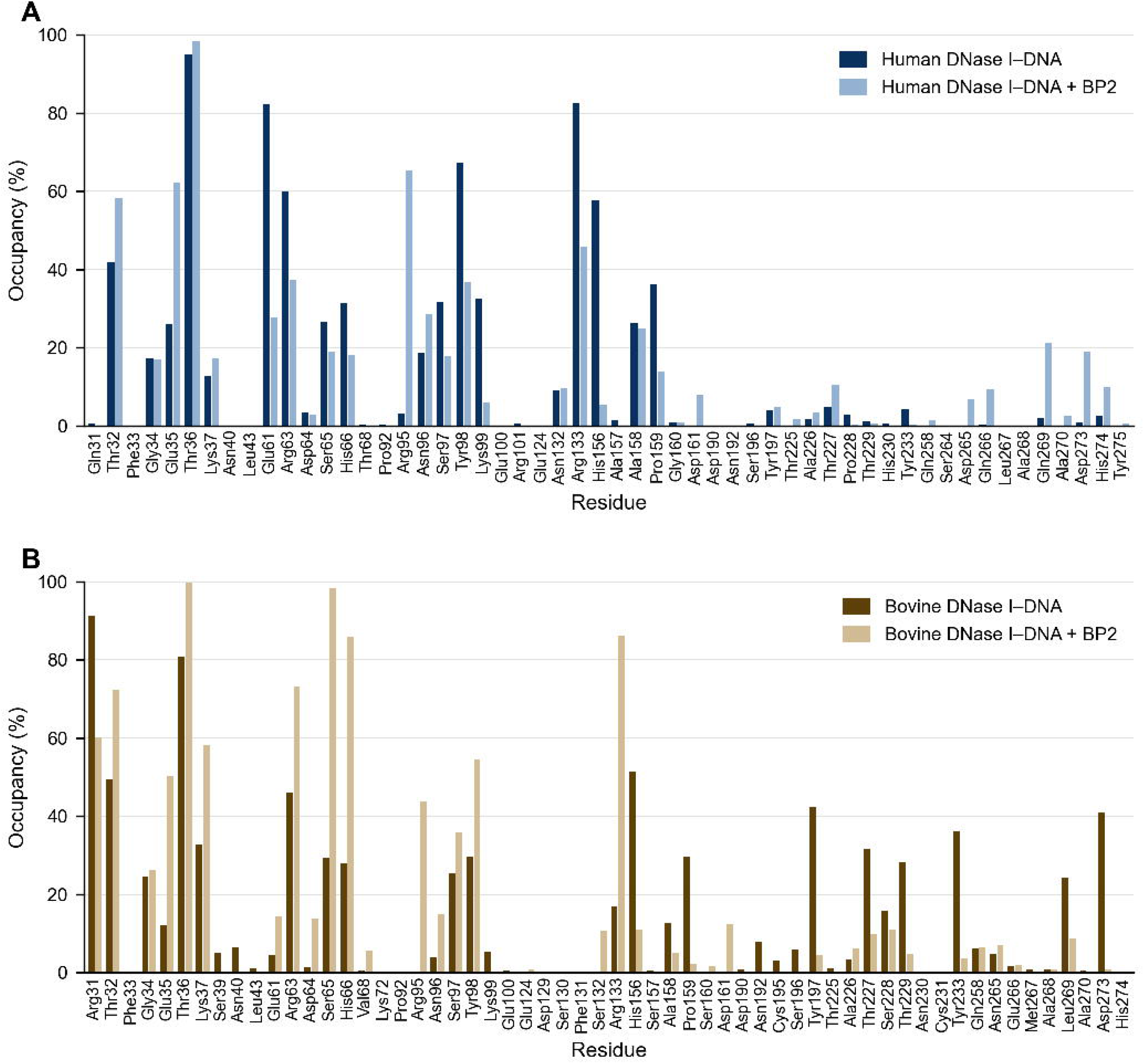
Effect of BP2 on DNA binding by DNase I in molecular dynamics simulations. Per-residue DNA contact occupancy for (A) human and (B) bovine DNase I during 100 ns simulations of the DNase I–DNA complex without (dark bars) and with (light bars) BP2. Occupancy is the percentage of trajectory frames in which a residue forms any interaction with the DNA. Residues are numbered according to the full-length UniProt sequences (human P24855, bovine P00639); subtract 22 for mature-protein numbering.

The presence of BP2 substantially reorganized the DNA contacts. The most notable effect was at the catalytic His156, whose contact with DNA fell from 58% to 5% in human DNase I and from 52% to 11% in bovine DNase I. His156 is conserved in 99.5% of the DNase I orthologs analyzed (Figure 2), and its loss of DNA contact in the presence of BP2 was observed in both the human and bovine simulations. Conversely, Glu35 gained contact in both species (from 26% to 62% in human and from 12% to 50% in bovine DNase I), and Arg95, which is not among the DNA-binding residues described in the crystal structures, established a new contact (from 3% to 65% in human and from 0% to 44% in bovine DNase I). Other changes were species-specific: Arg63, Tyr98, and Arg133 lost DNA contacts in human DNase I but gained them in bovine DNase I, where the DNA also shifted away from Tyr197, Thr229, and Tyr233 toward Ser65 and His66 (Figure 5).

Together, these simulations suggest that BP2 does not prevent DNA binding but instead disrupts the network of contacts that positions the DNA at the catalytic site, primarily at His156. This displacement of the DNA from the catalytic histidine supports the hypothesis that BP2 may act as a DNase I inhibitor.

### Experimental validation

To experimentally validate the computational predictions, we performed time-course inhibition assays of DNase activity in the presence of PDCs on a suspension of commercial bovine and human DNAse I. DNA hydrolysis was monitored by measuring the decay of normalized over time in the absence or presence of BP2, from which the initial rates were calculated. In the assays with the bovine enzyme, the presence of BP2 at a final concentration of 10⁻⁵ M reduced the initial rate by 21.5% compared with the control (p < 0.01, Figure 6). The same assay was repeated with recombinant human DNase I: the addition of BP2 at 10⁻⁵ M reduced the initial rate by 23.2% (p < 0.001, Figure 6). These changes in protein function in both species are consistent with the in silico observations and show that the impact previously described for zebrafish correlates with effects in other terrestrial species that may be exposed to the same contaminants.

**Figure 6.**
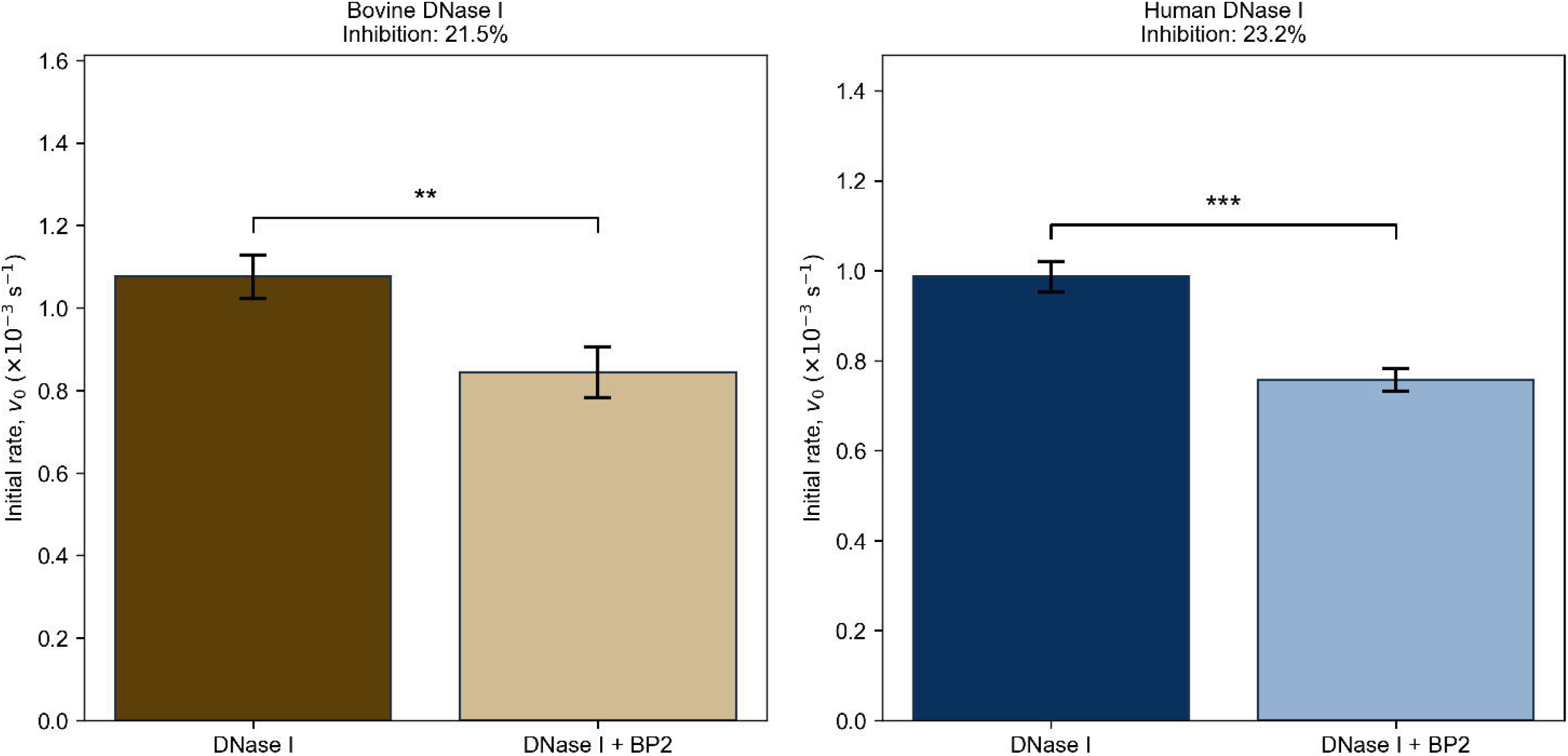
Inhibitory effect of BP2 on DNase I activity. Initial rates of DNA hydrolysis (v₀, calculated as the rate of decrease of buffer-subtracted, normalized fluorescence, –dF/dt, in ×10⁻³ s⁻¹) are shown for the control and in the presence of 10⁻⁵ M BP2, for bovine DNase I (left; P00639) and recombinant human DNase I (right; Pulmozyme, Roche; P24855). Bars represent the mean ± SD. Results were obtained from at least 5 independent experiments in all cases, with a total of n = 22–30 replicates per condition. For each experiment, v₀ was calculated as the slope over a common time window spanning the most linear sections (window of maximum R²) of all its replicates. Inhibition was calculated as 1 − (mean v₀ with BP2 / mean v₀ control). Differences between conditions were evaluated with Student’s t-test.

To rule out effects arising from simple sequestration of free Mg²⁺ cofactor by the compound, the assay was repeated with twice the MgCl₂ concentration in the reaction buffer (from 2.5 mM to 5 mM). Under these conditions, inhibition by BP2 persisted, indicating that the observed effect is not due to limited availability of Mg²⁺. This result is consistent with BP2 interacting with the protein, in line with the in silico prediction, rather than acting as a simple chelating agent of divalent cations.

### Conclusion

We established a computational strategy to identify potential human molecular targets of environmental pollutants. By integrating transcriptomic data with structural bioinformatics, we developed a reproducible, scalable pipeline that prioritizes human proteins for interaction with plastic-derived compounds. Docking predictions identified six targets for which PDCs outperformed the canonical ligand in binding affinity. These six candidates represent high-confidence hits suitable for experimental validation and for potential development as biomarkers of pollutant exposure.

By analyzing protein–ligand contact patterns and chemical similarity, we determined that PDCs do not act as structural mimics of endogenous ligands. Instead, interaction fingerprint analysis showed that PDCs adopt distinct binding modes predominantly driven by hydrophobic and π-type interactions. In addition, chemical similarity analysis using Morgan fingerprints yielded a median Tanimoto coefficient of only 0.054 between PDCs and their corresponding canonical ligands, confirming that these contaminants do not structurally resemble natural substrates. Together, these findings suggest that PDCs interfere with protein function not by directly competing for canonical chemical contacts, but by exploiting alternative interaction landscapes within the binding cavity, potentially inducing allosteric or conformational changes.

The functional profile of the prioritized candidates further underscores their toxicological relevance. Gene Ontology enrichment analysis showed that proteins with preferential affinity for PDCs are significantly enriched in immune response, programmed cell death, and extracellular matrix organization, as well as hydrolase and GTPase activities. This molecular signature closely aligns with the established toxicology of phthalates and bisphenols, which are widely reported as endocrine disruptors and modulators of inflammatory and apoptotic pathways. The convergence between our computational predictions and published toxicological data provides strong external validation for the biological plausibility of our hits. A relevant biological case among the putative targets is DNase I, which is inhibited by BP2. This enzyme plays a critical role in clearing neutrophil extracellular traps (NETs) and degrading extracellular DNA released during apoptosis; reduced DNase I activity has been experimentally linked to the development of systemic lupus erythematosus. Molecular docking revealed that BP2 binds within the enzyme’s active site with substantially stronger predicted affinity than the canonical ligand PO₄³⁻ (median binding energy −5.85 kcal/mol vs −2.55 kcal/mol). Strikingly, this binding mode is conserved in both bovine and human DNase I, supported by a highly similar binding-site structure (RMSD < 2.5 Å) that exhibits comparable affinities for the contaminants. Extending this conserved interaction to a livestock species suggests that BP2-mediated inhibition may affect animal health as well as human health via the food chain. We experimentally validated these predictions using in vitro activity assays, confirming that 10⁻⁵ M BP2 reduces the initial DNA hydrolysis rate by 21.5% in bovine DNase I (p = 0.007) and by 23.2% in recombinant human DNase I (p < 0.001).

Beyond confirming a single prediction, the *in vitro* assays show that the pipeline yields hits that withstand experimental scrutiny: BP2 inhibits DNA hydrolysis by both human and bovine DNase I, the effect persists when Mg²⁺ is supplied in excess –ruling out cofactor sequestration–and its reproducibility across species follows directly from the cavity conservation predicted in silico. This is particularly relevant for BP2, a compound whose regulatory status remains unsettled: the European Scientific Committee on Consumer Safety recently concluded that it could not establish its safety in cosmetic products, citing unresolved indications of genotoxicity, demonstrated endocrine activity, and missing repeated-dose and reproductive toxicity data ^36^. Identifying a conserved human target for such a compound turns an open regulatory question into a tractable molecular hypothesis, and illustrates the kind of mechanistic entry point the pipeline can offer for contaminants that still lack comprehensive toxicological assessment.

In conclusion, our work presents a transferable framework that bridges environmental toxicology and structural bioinformatics to uncover non-mimetic protein targets of chemical pollutants without requiring prior experimental binding data.

## Materials and Methods

### Proteins targets recruitment

The starting set of genes was obtained from a published transcriptomic study of zebrafish exposed to plastics ^22^. Zebrafish gene IDs were mapped to human orthologs using the Ensembl database ^34^, yielding 159 human orthologous proteins. From this set, we retained only those proteins for which a canonical ligand could be identified in the Protein Data Bank (PDB) ^35^, resulting in 27 human proteins.

However, these structures exhibited an average of 12% missing residues and lacked co-crystallized canonical ligands, making them unsuitable for docking. Therefore, we retrieved high-confidence structural models from the AlphaFold database (Hallgren et al. 2022), retaining the best-ranked model for each protein (pLDDT 65.5–97.3; 25 of 27 above 80). All subsequent docking analyses were performed on these AlphaFold models ^36^.

### Ligands preparation

A set of 12 plastic-derived compounds (PDCs) frequently reported in toxicological studies was initially compiled ^37,38^. SMILES notations were obtained from PubChem ^39^, and three-dimensional structures were generated using RDKit. To reduce the number of compounds for docking, we clustered them by chemical and structural similarity using the Butina algorithm, which yielded eight clusters. One representative compound per cluster (the centroid) was selected as the final PDC set: five phthalates (BBP, DPP, DEHP, DIOP, DTDP) and three phenolic compounds (BP2, BP3, BPA). Physicochemical descriptors (molecular weight, LogP, TPSA, HBD, HBA, rotatable bonds, aromatic rings, Fsp³ fraction, and QED) were calculated for each selected PDC using RDKit.

Canonical ligands for each target protein were identified as the co-crystallized molecules in the PDB entry corresponding to the protein’s biological substrate or known physiological ligand. To distinguish biologically relevant ligands from crystallization artifacts, we excluded common solvents and buffers using a previously published list ^40,41^.

### Binding sites assignments and Molecular docking performing

To facilitate comparison between canonical ligands and ligands of interest, we considered only the cavities predicted with CaviDB ^42^ that overlapped with annotated canonical ligand-binding sites in UniProt. To establish this correspondence, we calculated the fraction of residues in the predicted cavity that are part of the ligand-binding site, using a minimum overlap threshold requiring that at least 30% of the residues in the predicted cavity match residues of the binding site. In cases with multiple assignments for the same ligand, the one with the highest precision was selected.

The software chosen for molecular docking was AutoDock Vina ^43^. Ligand and receptor preparation followed the steps described in its manual. For each molecular docking run, a search box delimiting the region where the process takes place was defined. The center of the box was placed at the geometric centroid of the cavity atoms predicted by CaviDB. The box dimensions were calculated based on the size of the cavity, the target ligand, and, when available, the canonical ligand (exhaustiveness = 16). This procedure was repeated for each protein in the dataset.

Protein-ligand affinity estimates were performed by molecular docking using AutoDock Vina. Given that Vina has a standard error of 2.85 kcal/mol, this value was used as a threshold to determine whether one ligand has a better binding energy than another.

### Selection of species for analysis of the impact of plastic derivatives on the health of the ecological niche

From a food perspective, animals are the final stage of bioaccumulation before reaching the plate; phthalate esters (PAEs) incorporated by these animals through drinking water, contaminated forage, packaging materials, or handling materials can be transferred to their edible tissues and derived products such as milk, eggs, and meat, constituting a direct source of human exposure ^44^. For this reason, we considered it pertinent to recognize that a disruptor that alters the enzymatic activity of livestock may have health and productivity consequences that ultimately also affect human health, and therefore its study and consideration are relevant. Following the guidelines established by the European Food Safety Authority (EFSA), we included in the present study Bos taurus intended for meat production for human consumption ^45,46^.

### Evolutionary conservation analysis of DNase I

To evaluate the functional and structural relevance of the residues identified in the docking and transcriptomic analyses, we performed an evolutionary conservation analysis of DNase I. We retrieved 153 DNase I orthologous sequences from public databases and subjected them to multiple sequence alignment using standard alignment tools. The alignment was manually inspected and refined to ensure correct positional homology. We then estimated site-specific evolutionary rates using Rate4Site ^47^, which employs an empirical Bayesian framework to infer conservation scores based on the phylogenetic tree of the aligned sequences. The human DNase I sequence (UniProt ID: P24855) was used as the reference for residue numbering throughout the analysis. Evolutionary rates were reported on a scale from 1 (most variable) to 9 (most conserved), as defined by the Rate4Site algorithm. Residues with a rate score ≥ 7 were considered highly conserved, whereas those with scores ≤ 4 were considered variable. This approach allowed us to correlate sequence conservation with the known functional roles of catalytic, calcium-binding, and DNA-interacting residues previously characterized in the literature.

### In vitro experimental validation: DNase I activity assay monitored by fluorescence

DNase I activity was assessed by real-time monitoring of the decrease in fluorescence of Hoechst 33258 dye bound to DNA. This reagent markedly increases its fluorescence quantum yield upon binding to the minor groove of double-stranded DNA (preferentially A/T-rich regions), so progressive DNA fragmentation by the enzyme produces a sustained decrease in signal.

Reactions were carried out in a final volume of 100 µL in a microtitre plate and contained genomic DNA at a final concentration of 10 ng/µL and Hoechst 33258 at 1 µg/mL in reaction buffer (10 mM Tris-HCl pH 7.5, 2.5 mM MgCl₂, 0.5 mM CaCl₂, no EDTA). Before reaction initiation, the DNA–Hoechst 33258 mixture was pre-incubated for 20 min at 37 °C to allow formation of the DNA–dye complex. The reaction was initiated by adding bovine DNase I to a final concentration of 0.1 mU/µL (equivalent to 0.01 U per reaction).

For inhibition assays, DNase I was pre-incubated for 20 min at 37 °C in the presence of the inhibitor compound before addition to the DNA–Hoechst mixture. Compounds were prepared as 10⁻³ M stock solutions in PBS supplemented with 10% (v/v) ethanol, and 1 µL was added per well, yielding a final concentration of 10⁻⁵ M in the reaction. As a vehicle control, the same volume of solution without compound was added (final ethanol concentration 0.1% v/v). In preliminary validation assays, BP2 was used.

Measurements were performed on a plate reader in top-read mode, with excitation and emission wavelengths of 350 nm and 460 nm, respectively. Fluorescence was recorded every 15 seconds for 20 minutes at 37 °C. All experimental conditions were evaluated in triplicate.

Fluorescence values were corrected by subtracting the corresponding blank (reaction buffer supplemented with Hoechst 33258 in the absence of DNA) and then normalized to the initial value of each well to allow comparison between different experimental conditions. The initial reaction rate was determined from the slope of a linear regression over the first 500 seconds of the fluorescence curve, an interval in which the signal shows linear behaviour. The rate was expressed as the negative derivative of fluorescence with respect to time (−dF/dt), corresponding to the rate of fluorescence decrease, in units of s⁻¹.

To evaluate whether the observed inhibition depended on divalent cofactor availability, the assay was repeated with double the concentration of MgCl₂ in the buffer (5 mM MgCl₂), while keeping the remaining conditions constant.

### In silico validation: DNase I molecular dynamics

To investigate the structural basis of modulation by plastics, we carried out a detailed structural analysis using molecular dynamics simulations based on the complexes obtained in the previous step. Molecular dynamics simulations were performed with GROMACS ^48^ to explore structural changes in selected biological targets in the presence of plastic molecules. Additionally, a simulation run was performed with the canonical ligand. Each simulation was run for 100 nanoseconds using an all-atom representation with the CHARMM36 force field and the TIP3P water model ^49^. The v-rescale thermostat was used to maintain the temperature (T = 310 K) with a relaxation time of 0.1 ps, and Berendsen pressure coupling (1 atm) with a relaxation time of 1 ps was applied in the NPT ensemble . Pressure was coupled isotropically in the XY plane and the Z direction. The time step was 2 fs. The last frame of this equilibrated system was used as the initial configuration for a second simulation. The Particle Mesh Ewald (PME) method was used to treat electrostatic interactions ^50^, with a real space cutoff of 4 nm, a grid spacing of 0.16 nm, and cubic interpolation. We observed that 100 ns simulations are enough for dynamically calculating the binding contacts. To assess whether BP2 affects DNA binding, for each protein residue, we computed the percentage of simulation frames in which it contacted the DNA. In the absence of BP2, DNA was held mainly by residues previously described as DNA-binding in the DNase I-DNA crystal structures.

## Authors’ contributions

AH, LS, and AJVR carried out the bioinformatic analysis. LRS carried out the experimental analysis. AJVR and MOF carried out the health and ecological analysis and conceptualization. AJVR conceived the study and was in charge of its design and coordination. GP carried out the project administration and funding acquisition. All authors participated in drafting the manuscript and in its final reading and approval.

## Acknowledgements

LMS, AJVR, LRS and MOF are researchers from Consejo Nacional de Investigaciones Científicas y Técnicas (CONICET). This work was supported by Universidad Nacional de Quilmes (PUNQ 918/22). This work used computational resources from UNC Supercómputo (CCAD) – Universidad Nacional de Córdoba (https://supercomputo.unc.edu.ar), which are part of SNCAD, República Argentina. IBYME, where MOF has her lab, is supported by: Fund. René Barón, Fund. Williams and Fund Honorio Bigand.

## Data availability

All data can be accessed at the CONICET open data repository: https://hdl.handle.net/11336/301419. Additional data will be shared upon request.

