## supplementary_table for "In silico pipeline for identification of targets of plastic-derived chemicals"

### Supplementary Material

**Supplementary Table 1. KEGG disease enrichment analysis of proteins showing preferential binding affinity for plastic-derived compounds (PDCs) over their canonical ligands.** Only diseases with at least one target protein meeting the binding preference criteria are shown.

| KEGG Disease ID | Disease Name | Target Protein(s) | PDC Preference | Mechanistic Relevance |
|---|---|---|---|---|
| H00080 | Systemic lupus erythematosus | DNase I (P24855) | BP2, BP3, BPA | DNase I inhibition impairs NET clearance |
| H00211 | Hemochromatosis | Ferritin (P02794) | BP2, BPA, BBP, DEHP | Ferritin dysregulation in iron overload |
| H00079 | Asthma | ALOX5 (P09917) | BP2, BP3, BPA, BBP | Leukotriene synthesis pathway |
| H00276 | Galactosialidosis | Cathepsin L2 | BP2, BP3, BPA | Lysosomal dysfunction |
| H01742 | Coronary artery disease | ALOX5, FABP7, ACBP | Variable | Lipid metabolism and inflammation |
